# Development of a sensitive molecular diagnostic assay for detecting *Borrelia burgdorferi* DNA from blood of Lyme disease patients by digital PCR

**DOI:** 10.1101/2020.06.16.154336

**Authors:** Srirupa Das, Denise Hammond-McKibben, Donna Guralski, Sandra Lobo, Paul N. Fiedler

## Abstract

Lyme disease patients would benefit greatly from a timely, sensitive and specific molecular diagnostic test that can detect the causal agent, *Borrelia burgdorferi*, at the onset of symptoms. Currently available diagnostic methods recommended by the Centers for Disease Control and Prevention for Lyme disease, involve indirect serological tests that rely on the detection of a host-antibody response which often takes more than three weeks to develop. This results in non-detection of many genuine cases on a timely basis, preventing complete cure. In this study we have developed a digital PCR (polymerase chain reaction) assay that detects Lyme disease on clinical presentation at twice the sensitivity of the currently available diagnostic methods, using a cohort of patient samples collected from the Lyme disease endemic state of Connecticut, USA in 2016-2018. Digital PCR technology was chosen as it is more advanced and sensitive than other PCR techniques in detecting rare targets and the lower limit of detection of this diagnostic assay was found to be three genome copies of *B. burgdorferi*. The paucity of spirochetes in the bloodstream of Lyme disease patients that hinders the clinical adoption of PCR-based diagnostic tests, was overcome by using a comparatively larger sample volume, pre-analytical processing of blood samples and a pre-amplification step to enrich for *B. burgdorferi*-specific gene targets before using the digital PCR technology to analyze patient samples. Pre-analytical processing of blood samples from acute patients revealed that the best sample type for Lyme disease detection is platelet-rich plasma and not whole blood. If detected on time, Lyme disease can be cured completely limiting the overuse of antibiotics and associated morbidities.

## Introduction

Lyme disease (LD), a systemic tick-borne infection caused by the bacteria *Borrelia burgdorferi*, is the most common vector-borne disease in the USA. According to the Centers for Disease Control and Prevention (CDC) there is an estimated 300,000 new cases of LD in the USA every year. However, only 10% of these cases are actually reported and diagnosed [1]. Due to the non-specific flu-like symptoms of LD and the lack of any reliable testing during the early stages of infection, diagnosis becomes very challenging. According to the CDC, the characteristic symptom of LD-a typical bulls-eye rash known as Erythema Migrans (EM) develops in only 70-80% of the patients and can often be confused with other similar rashes [2]. The current CDC approved diagnostic methods to detect LD is serological two-tiered testing (TT testing), which includes a screening test by ELISA and a specificity test by western blot. This test is inaccurate in the early stages of disease as it relies on the indirect detection of a host antibody response that often takes three weeks or more to develop. As a result 25-50% of positive LD cases are missed during initial diagnosis. The CDC cautions that because the test is not likely to be positive until 3-6 weeks post-infection, doctors who suspect LD based on symptoms and epidemiological information, should prescribe antibiotics even if the test is negative [3]. Early diagnosis is critical to minimize the long term effects and morbidity associated with LD and ensure complete cure.

Methods to directly detect LD causing bacteria by culturing and/or polymerase chain reaction (PCR) have not been much successful till date. While *B. burgdorferi* is recalcitrant to culturing under laboratory conditions, a clinically relevant PCR assay for LD detection from blood has not been established due to the insufficient sensitivity of conventional PCR methods and the extremely low levels of the spirochete found in the blood of infected patients [4, 5, 6, 7]. In the past, the use of PCR methods to detect *Borrelia* infection from blood of acute LD patients suffered from low sensitivities of around 18.4% and 26.1% [8, 9]. As a result, the limit of detection of these assays is below the threshold necessary for reliable LD detection in clinical blood samples. The PCR results were also discordant depending on the type of specimens tested and the symptoms that the patients reported, limiting the clinical adoption of PCR testing for LD [4, 5, 6, 7]. Current advances in molecular techniques have led to better DNA extraction and amplification techniques, resulting in detection of low copy numbers of *Borrelia* DNA from larger volumes of patient samples [10]. Prior culturing of *B. burgdorferi* under laboratory conditions from patient samples followed by PCR has resulted in better detection rates, indicating the unusually low bacterial load in humans [11]. Adoption of newer PCR techniques like quantitative real-time PCR (qPCR) and nested PCR in LD detection has demonstrated some improvement in the sensitivity of detection of *B. burgdorferi* by PCR [11, 12, 13]. Low bacterial load of the spirochete in circulating blood of infected humans, have made investigators use a larger volume of patient blood to boost the detection rates of LD [7]. In recent times, nanotrap technology has been applied to detect the outer surface protein A of *B. burgdorferi* that is shed in the urine of patients afflicted with LD [14].

Development of a very sensitive diagnostic test to directly measure the *B. burgdorferi* in blood would significantly enhance the detection of LD in the early stages, when treatment is most effective. In this study we have used digital PCR (dPCR) to develop a sensitive method to detect LD directly at clinical presentation. Digital PCR is a type of quantitative PCR method that provides a sensitive and reproducible way of measuring the amount of DNA or RNA present in a sample. During dPCR, the initial sample mix is partitioned into a large number of individual wells prior to the amplification step, resulting in either 1 or 0 targets being present in each well.

Following the PCR amplification, the number of positive versus negative reactions is determined and the absolute quantification of target is calculated using Poisson statistics [15]. As compared to other PCR methods, the partitioning of samples during dPCR leads to a significant improvement in the sensitivity of the technique to detect rare alleles, low pathogen load and targets in limited clinical samples [16]. In order to further improve LD detection in patients, we have also incorporated a pre-analytical processing step of blood samples and a pre-amplification step to enrich for *B. burgdorferi*-specific target DNA, prior to analyzing them by dPCR. The assay we developed can detect LD at twice the sensitivity of the current CDC-recommended diagnostic methods.

## Materials and Methods

### Ethics Statement

This study was approved by Biomedical Research Alliance of New York Institutional Review Board. The participants provided us with written informed consent prior to inclusion in the study.

### Culture of *B. burgdorferi* strain

*Borrelia burgdorferi* strain, B31 was purchased from ATCC, Manassas, Virginia (Catalogue No. #35210) and maintained in complete BSK-H media, which included 6% rabbit serum (complete media from Sigma-Aldrich, St. Louis, Missouri) at 33^°^C.

### Collection of LD patient samples

Paired whole blood (WB) and serum samples were collected from 46 clinically diagnosed LD patients during 2016-2018 from a LD endemic area (Connecticut, USA). During the course of the study, seven of these patients dropped out after the initial visit and three patients dropped out after the second visit. Samples were collected from each patient at the initial pre-treatment (acute), during treatment (2 weeks post-diagnosis) and post-treatment stages (6 weeks post-diagnosis). Patients were referred to the study by their primary care provider immediately following their LD diagnosis. Patients included in the study most often presented with a rash consistent with EM, a known tick-bite, fever and other symptoms consistent with *B. burgdorferi* infection. Patients who had a known history of LD during the past 5 years, who were pregnant or those who had been taking antibiotics for more than 72 hours were excluded from the study. WB from patients was collected into Cyto-Chex® BCT tubes (Streck, La Vista, Nebraska) and a Vacutainer SST tube (BD Biosciences, San Jose, California). Non-LD controls were collected from the states of Connecticut, USA (100 samples) under our approved IRB and from Tennessee, USA (30 samples purchased from Tennessee Blood Services, Memphis, Tennessee) We also obtained de-identified blood samples from clinically diagnosed LD patients with positive IgM western blot from Danbury Hospital, Danbury, Connecticut under our approved IRB protocol for optimization of the assay.

### Serological analysis

Serum samples were subjected to TT-testing for *B. burgdorferi* at Danbury Hospital, following recommended guidelines of the CDC [17]. Serum samples were also sent to the Mayo Clinic, Rochester, Minnesota for C6 peptide Lyme ELISA testing. The TT-testing results (supplemental **S1 Table**) were used to compare the efficiency of the Lyme PCR diagnostic assay that we have developed.

### Pre-analytical processing of blood samples

The WB samples were subjected to pre-analytical processing before they underwent DNA extraction. The blood samples were processed in such a way so that we could save them as serum, plasma, Platelet-Rich-Plasma (PRP) and WB per patient. All the samples were aliquoted @ 1ml per cryo-vial and stored at -80°C before use. For serum collection, the Vacutainer SST tube was centrifuged at 1000 g for 10 minutes at 4°C and the supernatant was collected. Plasma was collected by centrifuging the whole blood at 1200g for 10 minutes at 4°C and the supernatant was stored. For PRP collection, the whole blood was centrifuged at 260g for 10 minutes at room temperature (RT) and the supernatant (above the buffy coat) was collected. However, the PRP from patient samples of 2017 were processed differently to test for the efficiency of different pre-analytical processing and storage methods. In 2017 the PRP was pelleted down by another round of centrifugation at 15,000rpm for 10 minutes at 4°C. The supernatant was discarded and the PRP pellet was stored at -80°C till further use.

### DNA extraction and precipitation

DNA extraction was done from 1 ml of the different sample types using QIAamp DNA mini kit (Qiagen, Hilden, Germany), as per the vendor’s instructions with slight modifications. All the samples were pelleted down at 15,000rpm for 10 minutes at 4°C and the supernatant was discarded. The pellets underwent bacterial DNA isolation by re-suspending in 180µl of Buffer ATL and 20µl of Proteinase K. The lysate underwent 1 hour of incubation at 56°C in a thermomixer (Eppendorf, Hamburg, Germany) with shaking. Following this, 200µl of Buffer AL was added along with 10µg of poly (A) carrier DNA (Roche, Basel, Switzerland) to the lysate and incubated for an additional 10 minutes at 70°C. Then 230µl of molecular-biology grade ethanol (Sigma-Aldrich, St. Louis, Missouri) was added to the lysate before passing through a QIAamp mini spin column at 6000g for 1 minute at RT. The filtrate was discarded and the spin column was washed with wash buffers AW1 and AW2, as per the vendor’s instructions. Finally DNA was eluted with 150µl of Buffer AE (pre-heated to 65°C) twice resulting in a total volume of ∼300µl. Bacterial DNA extraction from whole blood was done with QIAamp DNA blood mini kit (Qiagen, Hilden, Germany) following vendor recommended protocols.

### Processing of extracted DNA

The total volume of extracted DNA was precipitated with 1/10^th^ volume of 3M sodium acetate (Invitrogen, Carlsbad, California) and double volume of chilled molecular biology grade ethanol (Sigma-Aldrich, St. Louis, Missouri) at -20°C overnight. The precipitated DNA was pelleted down at 15,000rpm for 10 minutes at 4°C. The supernatant was discarded and the pellet washed twice with 700 µl of 70% ethanol by tapping and dislodging the pellet followed by centrifugation at 15,000rpm for 10 minutes at 4°C. The clean DNA pellet was dried in a sterile environment at 37°C for 1 hour or until dry. The DNA pellet was dissolved in 3.125µl/12.5µl of DNA suspension buffer (Teknova, Hollister, California) according to need. The re-suspended DNA was stored at -20°C until further use.

### *B. burgdorferi*-specific TaqMan assays

Bio-informatic tools (BLAST and Primer BLAST (NCBI); Multiple Sequence Alignment by CLUSTALW) were used to identify four unique *B. burgdorferi*-specific gene sequences for custom manufacturing of TaqMan assays (**Table 1**) by Life-Technologies (Carlsbad, California). *B. burgdorferi* strain B31 was the source of all sequences chosen from the four different genes: *ospA* (GenBank: AE000790.2), *ospC* (GenBank: U01894.1), *fla* (GenBank: X15661.1) and *rpoB* (GenBank: AE000783.1).

**Table 1.**
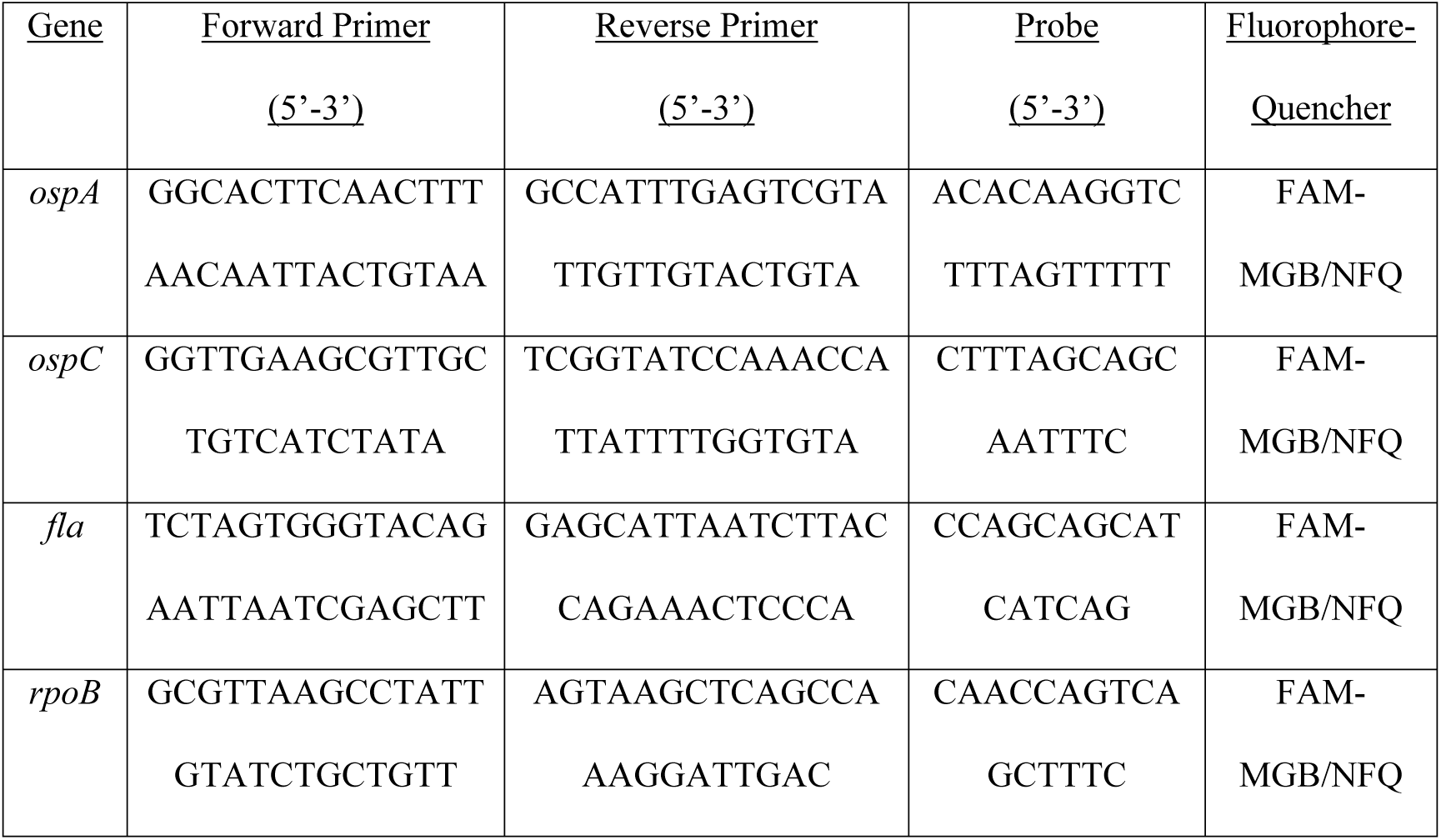
Primer and probe sequences of the TaqMan assays.

### Pre-amplification of DNA

*B. burgdorferi*-specific gene targets were enriched by pre-amplification PCR with DNA extracted from 1ml of sample. Total extracted DNA (3.125µl) was mixed with 6.25µl of TaqMan PreAmp Master Mix (Life-Technologies, Carlsbad, CA) and 3.125µl of TaqMan assay pool (made by diluting 100-fold of the TaqMan assays with nuclease-free water) for PCR at 95°C-10 minutes; 14 cycles of 95°C-15 sec, 60°C-4 min and a final hold at 4°C. Pre-amplified DNA was diluted 5-fold with DNA suspension buffer before use in PCR. In 2016 the extracted DNA was suspended in 12.5µl of DNA suspension buffer and only one-fourth (3.125µl) of the material was placed through the pre-amplification PCR. The rest of the procedure remained same.

### Molecular detection of *B. burgdorferi* genes from patient samples by dPCR and qPCR

Digital PCR was performed on BioMark platform (Fluidigm Corporation, San Francisco, California) with qdPCR37K Integrated Fluidic Chips, following the manufacturer’s instructions with slight modifications. Instead of using 1.8µl of DNA template, 2.1µl of the diluted pre-amplified DNA was used in each 6µl PCR reaction. Patient samples underwent dPCR with each of the four TaqMan assays in a singleplex format with TaqMan Gene Expression Master mix (Life-Technologies, Carlsbad, California) at: 50°C-2 minutes; 95°C-10 minutes; followed by 40 cycles of 95°C-15 sec and 60°C-1 min. Presence of a single red spot with sigmoidal amplification curve and a Ct value ≤23 in each panel was considered to be positive. QPCR was performed with 3µl of the diluted pre-amplified DNA as template and TaqMan Gene Expression Master mix in a total volume of 20µl, following the vendor’s instructions on a QuantStudio 7 Flex Real Time PCR System (Life Technologies, Carlsbad, California). The PCR parameters were as mentioned above. A Ct value of less than 35 was considered to be positive for detection. All PCR reactions were done in triplicates and repeated thrice to monitor reproducibility.

## Results

### Analytical specificity and sensitivity of *B. burgdorferi*-specific TaqMan assays

Specificity testing of TaqMan assays for the *B. burgdorferi*-specific genes (*ospA, ospC, fla, rpoB*) was performed with 1ng of total DNA from different sources (human, *Treponema denticola* and 27 clinically relevant microbial species listed in supplemental **S2 Table**) by dPCR and qPCR. Quantitative genomic DNA from *B. burgdorferi* (ATCC^®^ 35210DQ™) was used as positive control. None of the TaqMan assays showed cross-reactivity with any of the other pathogenic bacterial or human DNA (data not shown). Sensitivity of each TaqMan assay in detecting *B. burgdorferi* DNA was measured twice, once using blood spiked-in with cultured bacteria and repeated with *B. burgdorferi* quantitative genomic DNA. When analyzing blood spiked-in with cultured bacteria, it was found that all four *B. burgdorferi-*specific TaqMan assays were able to detect three *Borrelia* genome copies (data not shown). Since manual counting of spirochetes on a hemocytometer is approximate and subjective, we measured sensitivity of the TaqMan assays again using known amounts of *B. burgdorferi* quantitative DNA purchase from ATCC (Manassas, Virginia). The assay has a lower detection limit of three genome copies of *B. burgdorferi* for all the four genes tested in the dPCR format (**Fig 1A**). Due to statistical probability, when copy numbers of 1 or less were tested, we were able to detect *B. burgdorferi* DNA in some but not all panels. Similar results were obtained when qPCR was done to test the sensitivity of the TaqMan assays (**Fig 1B**).

**Fig 1.**
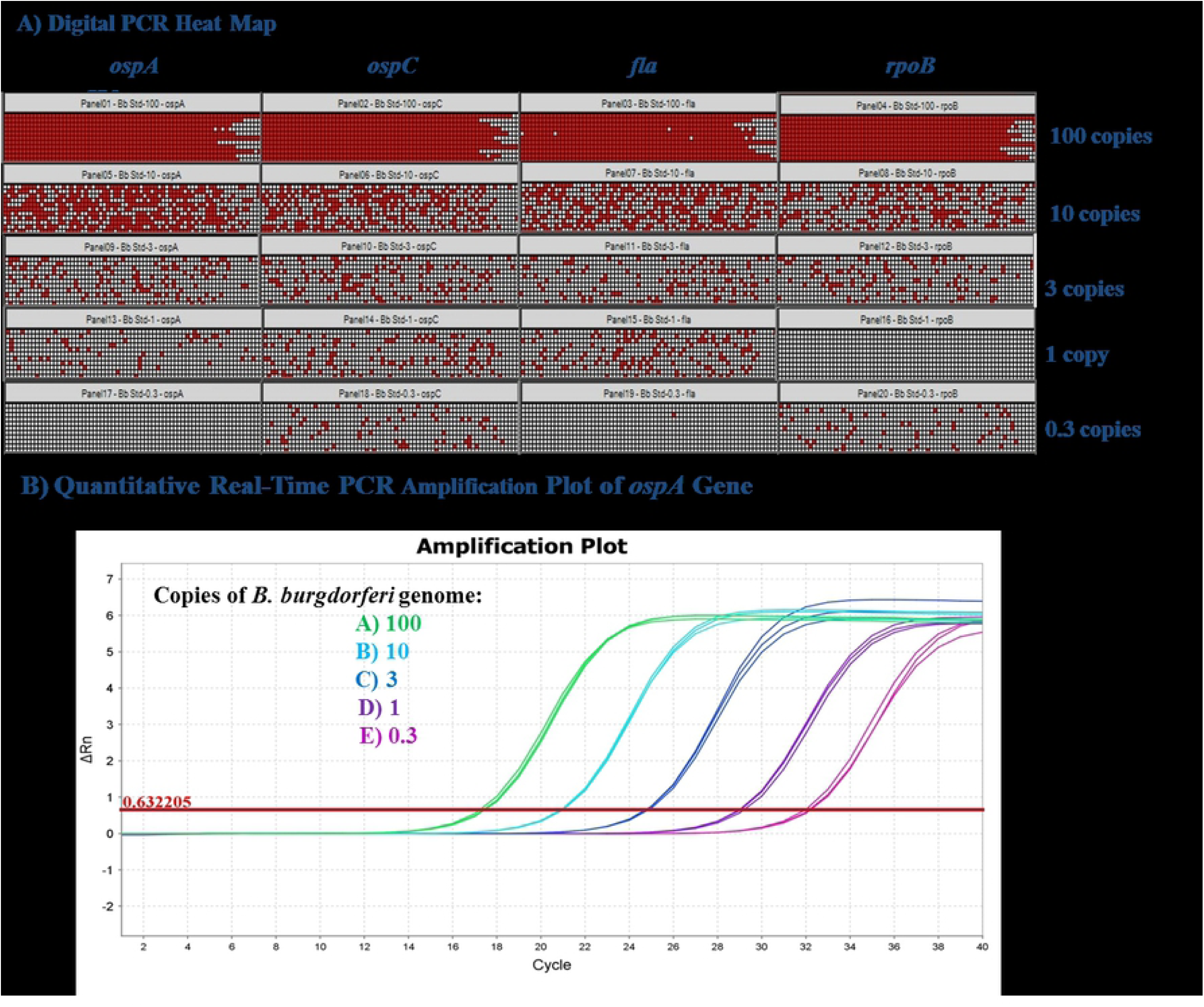
Sensitivity testing of *B. burgdorferi*-specific TaqMan assays. *B. burgdorferi* genomic DNA standard (ATCC) was diluted in poly (A) carrier DNA containing DNA suspension buffer and mixed with human DNA extracted from healthy control blood specimens. Serial dilutions were made to achieve 100, 10, 3, 1 and 0.3 genome copies and subjected to dPCR and qPCR with the *ospA, ospC, fla* and *rpoB* TaqMan assays. **(A)** Heat map depicting the detection of the four *B. burgdorferi* genes by dPCR and **(B)** representative amplification plot of qPCR with *ospA* TaqMan assay showing detection of low copy numbers of *B. burgdorferi* genome.

### Pre-analytical processing of blood samples

Pre-analytical processing of blood samples was optimized for *B. burgdorferi* detection using serum, plasma, PRP and WB spiked-in with 3 and 10 copies of cultured *B. burgdorferi*, separately. The sensitivity of detection for *B. burgdorferi* genes was the best when spiked-in PRP was used as a pre analytical sample source. Cultured bacteria were first spiked into the different matrices and then underwent the extraction procedure. The rate of detection of *B. burgdorferi* genes was very low when tested on spiked-in serum, plasma or WB, even when a pre-amplification step was included before the dPCR step. This indicated PRP to be the most effective sample type for detecting *B. burgdorferi* genes (data not shown). When clinically diagnosed LD IgM western blot positive samples from Danbury Hospital were used to test this hypothesis, the results re-confirmed that PRP was the most suitable sample type for detecting LD (**Fig 2**). In 2017 when patient samples collected under our approved IRB protocol were stored at -80°C before use as PRP pellet instead of PRP, the detection rate of *Borrelia* genes from PRP pellet decreased significantly. When PRP pellets of these 21 patients from their 3 serial visits (excluding the dropouts) were tested by qPCR, only four of the patients showed any significant Ct value for one of the four *B. burgdorferi* genes in our panel (supplemental **S3 Table**). A Ct (cycle threshold) value ≤35 was considered to be positive, to eliminate non-specific artifacts from consideration. Hence, the detection rate for LD of these 21 patients from PRP pellets was 19.05% after the three visits. The same results were obtained by dPCR (data not shown). These results indicate that variation in pre-analytical processing and storage as different sample types can have a drastic effect on the detection rate of *Borrelia* genes and PRP is the best sample type for detecting LD by either dPCR or qPCR.

**Fig 2.**
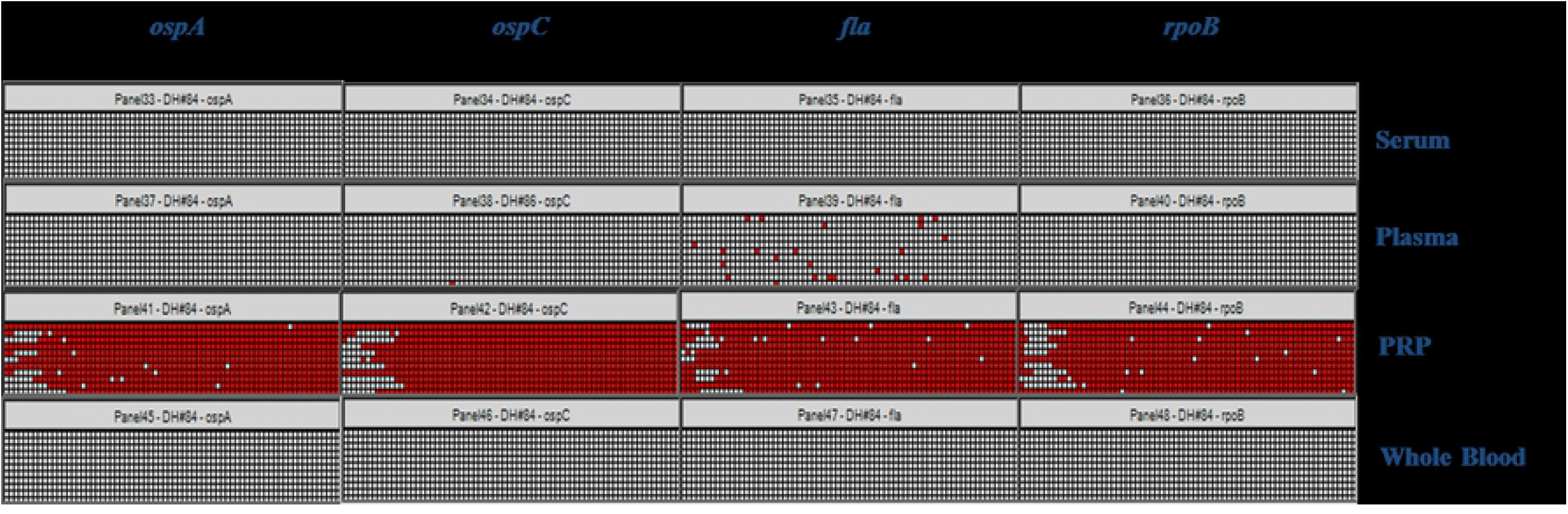
Determination of best sample type by pre-analytical processing of blood samples. Clinical blood samples which were positive by classical two-tier serology for Lyme disease were collected from Danbury Hospital, Connecticut and processed by different pre-analytical methods to get serum, plasma, PRP and whole blood, prior to DNA extraction. Following DNA extraction and pre-amplification to enrich for *Borrelia*-specific targets, the samples were subjected to digital PCR (dPCR) analysis for detection of *ospA, ospC, fla* and *rpoB* genes using TaqMan assays. PRP sample type was found to give the best sensitivity for detection of all four *B. burgdorferi* genes in the panel by dPCR.

### Evaluation of clinical samples with dPCR technology

After determining the best sample type to be used and optimization of the assay conditions, the patient samples collected under approved IRB protocol from 2016 and 2018 were subjected to dPCR assay to detect the four *B. burgdorferi*-specific genes. In 2016, serial blood draws were obtained from 11 patients of whom 2 patients dropped out after the initial visit. Patient samples were subjected to the steps of pre-analytical processing, DNA extraction, DNA precipitation and pre-amplification to enrich for *Borrelia*-specific gene targets followed by the optimized dPCR assay. Only one/fourth of the DNA extracted from 1ml PRP went into the pre-amplification step and analyzed by dPCR. Out of the 11 patients, dPCR assay could detect *Borrelia* DNA in 7 patients at either the initial or the 2 weeks post-diagnosis visits. There was a gradual clearing of the signal, as the duration of antibiotic treatment increased. A representative picture of the heat map from one clinical patient is shown in **Fig 3**. In 2018, we observed that 10 out of 14 patients during their initial visit were positive for at least one of the four *B. burgdorferi* genes in our panel. When compared to the classical TT-testing results, out of these 14 patients only 2 were detected as positive for LD by western blot during their initial visit (supplemental **S1 Table**). In 2018 modifications made to the pre-amplification protocol allowed us to analyze the total DNA extracted from 1ml of PRP by dPCR (instead of one/fourth of the DNA in 2016). However, in 2018 it was also observed all enrolled patients were positive for at least three of the four *B. burgdorferi* genes in our panel at 2 weeks and 6 weeks post-diagnosis stages. There was no evidence of much clearing of signal in these patients, as the duration of antibiotic treatment increased. When the dPCR data from 2016 and 2018 patient samples were compared against the classical TT-testing results, the sensitivity of our dPCR assay in detecting LD was at least two times more. At the initial visit, TT-testing showed positive results only in 24.35% of the cases compared to 58.54% detection rate of the dPCR assay (**Table 2**). We did not include the results of 2017 for calculating detection rates due to changes in patient sample processing and storage protocols which made a drastic difference in the detection rate of *B. burgdorferi* genes, as mentioned before. *B. burgdorferi* DNA was undetected in 100 Connecticut (endemic) and 30 Tennessee (non-endemic) blood samples with no symptoms of LD, validating our dPCR assay to be highly specific for detecting LD. A representative picture of endemic and non-endemic LD negative controls is shown in **Fig 4**.

**Table 2.** Comparison of test results for patients diagnosed with Lyme disease during the study.

| <u>Test Method</u> | <u>Percentage of positive samples(%)</u> |  |  |
| --- | --- | --- | --- |
|  | <u>At diagnosis</u> | <u>Two weeks Visit</u> | <u>Six weeks Visit</u> |
| Two-Tiered Serology | 24.35 | 45.84 | 43.33 |
| Digital PCR | 58.54 | 77.78 | 83.34 |

**Fig 3.**
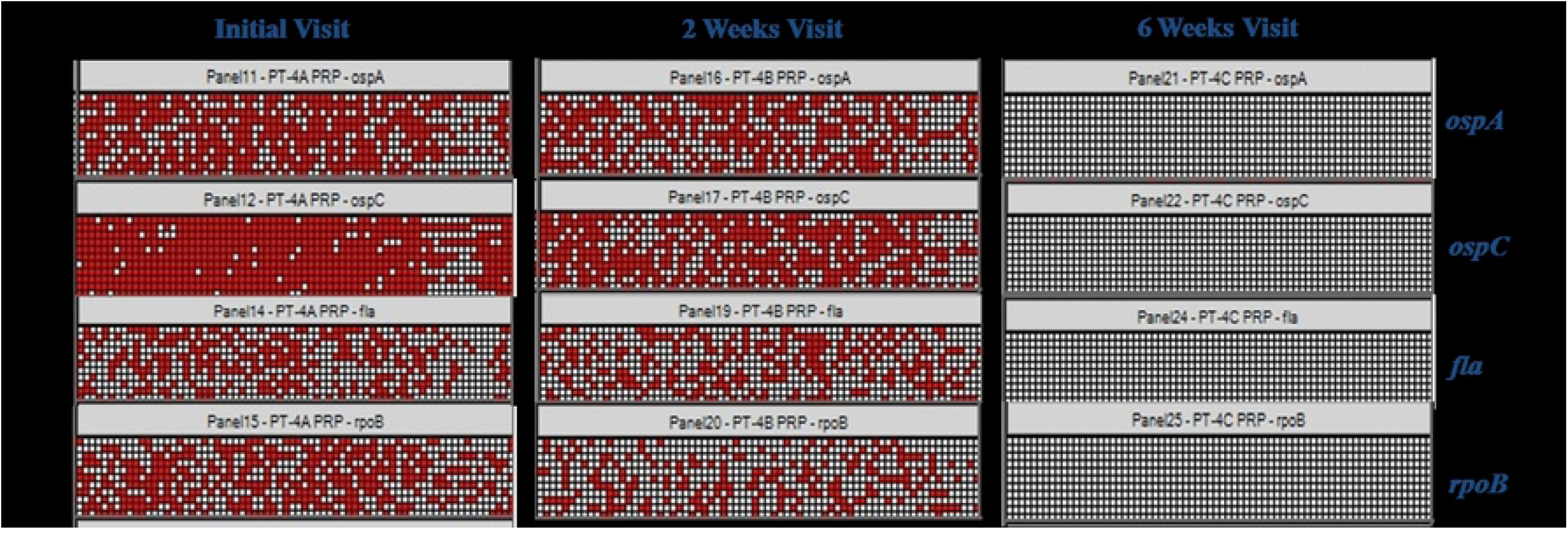
Analysis of Lyme disease patient samples by digital PCR. A representative picture showing the heat map of digital PCR analysis with the four *B. burgdorferi*-specific genes from one of the patients. The visits indicate the duration of antibiotic treatment.

**Fig 4.**
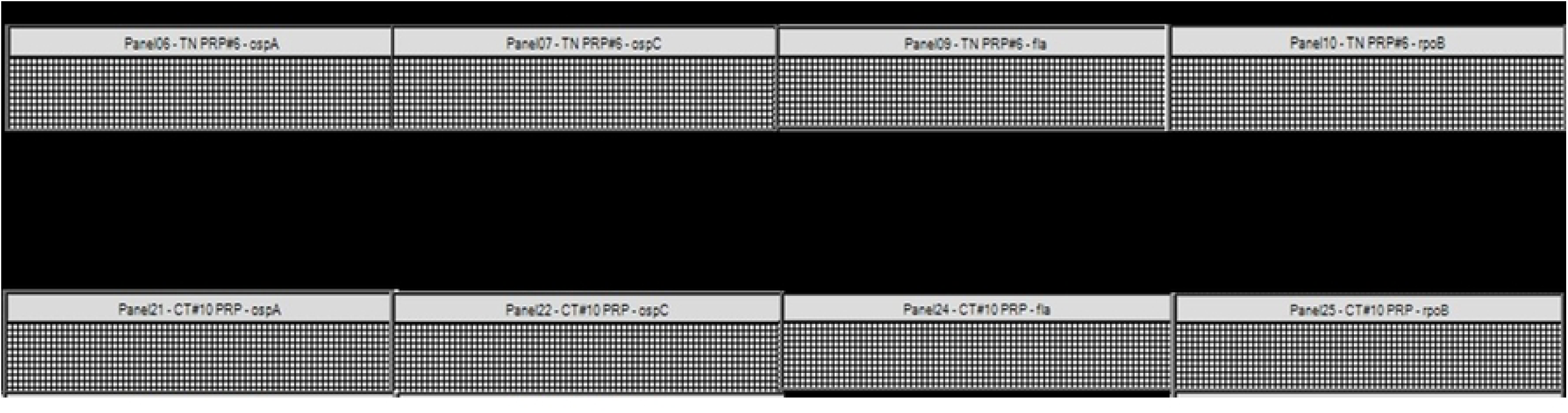
Digital PCR analysis of non-Lyme disease control patients. A representative picture showing the heat map of digital PCR analysis from **(A)** a Lyme endemic area negative control patient recruited from Connecticut and **(B)** a non-Lyme endemic area patient with no Lyme disease symptoms from Tennessee. No *B. burgdorferi* DNA was detected from the negative control patients.

## Discussion

Development of a highly sensitive and specific diagnostic method to detect LD during acute infection is critical to overcome the ineffectiveness of treatment when LD is detected in the late stages. The current CDC-approved TT-testing is insensitive due to the delayed antibody response during the first few weeks of infection and variations in the host immune responses. Moreover, the interpretation of TT-testing results is subjective resulting in false positive or false negative cases [18, 19]. Even the most characteristic symptom of LD, the EM rash is often either diffused or missing, preventing accurate diagnosis [20, 21]. This study has led to a much improved method to detect *B. burgdorferi* directly from the WB of patients in acute LD cases. Inclusion of various steps to improve sensitivity: pre-analytical processing, pre-amplification of *Borrelia-* specific gene targets prior to PCR and using the dPCR platform has led to the development of a robust assay to detect *B. burgdorferi* DNA from the blood of patients. By employing these methods, we were able to overcome the loss of assay sensitivity due to the presence of very low number of spirochetes in the blood stream of patients [11, 13, 22, 23]. The results from this study indicate that our dPCR assay has succeeded where previous assays involving PCR technology had failed to achieve the required sensitivity to become a clinically approved diagnostic assay for LD [8, 9].

This study was designed to recruit patients during the initial stages of infection, during the course of antibiotic treatment and after the completion of the antibiotic course. This cohort of patient samples formed the base on which this improved assay was developed. Adoption of dPCR technology achieved the highest sensitivity in detecting *B. burgdorferi* DNA and resulted in detection of as low as one to three genome copies of *B. burgdorferi* (**Fig 1**). Inclusion of a pre-analytical processing step to determine the best sample type for the detection of *B. burgdorferi* DNA made our assay more robust. This study clearly shows that PRP gives the best detection rates for *B. burgdorferi* genes, as compared to plasma, serum and WB (**Fig 2**). Presence of PCR inhibitors, such as hemoglobin, leukocyte DNA and IgG might have an impact on *Borrelia* DNA detection in WB and plasma [24] and also in all likelihood the spirochetes co-migrate in the PRP fraction. Recent studies have shown that better results are achieved when larger blood volumes are used in conjunction with other detection methods like nested PCR [11, 13] or isothermal amplification prior to multi-locus PCR/Electrospray Ionization Mass Spectrometry [4]. In our study we have used 1ml of PRP as the starting material to achieve high detection for the *Borrelia* genes. Depending on the hydration level of the patient, generally 1ml of PRP is obtained from 2ml of WB. This study has also revealed that the storage of the starting material has a huge impact on the sensitivity of the PCR assay. In 2017 when patient samples were stored at -80°C before use as PRP pellets instead of PRP, the detection rate of *Borrelia* genes from PRP pellets decreased significantly (supplemental **S3 Table**). Only 19.05% of the 21 clinical patients got detected for LD after the three serial visits.

Modifications made to sample processing and pre-amplification prior to dPCR increased assay sensitivity, overcoming the challenge of low circulating spirochete DNA in clinically diagnosed patient samples (**Fig 3**). By concentrating the DNA, we ensured that the maximum possible bacterial DNA amount was analyzed to improve the sensitivity of the assay. The pre-amplification step was essential to obtaining enhanced sensitivity in our assay, as *B. burgdorferi* DNA was not detected in samples that did not undergo this step (data not shown). Therefore, pre-amplification was included in the standard operating procedure to enrich for *Borrelia* DNA when analyzing patient samples. Due to assay modifications, the pre-amplification steps in 2016 and 2018 differed in the amount of DNA that was used in the analysis. In 2016, one-fourth of the extracted DNA from 1ml of patient sample went into the pre-amplification PCR, while in 2018 the entire extracted DNA was used. In spite of this discrepancy in the amount of DNA that was analyzed from patient samples in these two years, we achieved high detection rates of *Borrelia* genes (**Table 2**). A plausible explanation could be that patient recruitment in 2016 with stringent requirements (recruited patients definitely needed to have the EM rash for participation in the study) could have selected patients with a higher bacterial load and less sample variability justifying why the 2016 samples with less amount of analyzed DNA were detected like the 2018 patient samples (where the entire extracted DNA went into the pre-amplification step). In 2018, patients with or without the EM rash but meeting other criteria, as mentioned before in material and methods section, were included in the study. We did observe a gradual clearing of the signal as the antibiotic treatment continued in 2016 patient samples, as compared to 2018 patient samples. It is difficult to explain why this happened but in patient samples the integrity of the spirochetes is quite questionable: they can be fragmented, can form round bodies or blebs or even be hiding in biofilms [25, 26, 27]. This could lead to variability in extraction of spirochetal DNA from such patient samples using regular DNA extraction kits. Experiments evaluating extraction efficiencies of spirochete DNA (in its various forms) using modification in extractions buffers/methods/kits need to be carried for determining optimal methods.

Comparison of the dPCR and TT-testing results of the clinical patient samples revealed that the assay developed in this study was able to identify patients during the early stages of infection, when the immune response was yet to develop (**Table 2**). At clinical presentation, our assay was at least twice as effective in diagnosis when compared to the CDC-approved classical serology tests. Additionally, the dPCR assay is highly specific with no false positives, as none of the 130 negative controls got detected (**Fig 4**). The only limitation of this research work is the small patient pool that we had access to during the course of the study. A power calculation had revealed that 75 patients are required to achieve a statistically significant conclusion. Though more patient samples need to be tested by this assay, the result trends are promising and if applied to a clinical setting can lead to an early and accurate diagnosis of LD facilitating timely treatment, reducing the overuse of antibiotics and associated morbidities.

## Acknowledgements

The authors would like to thank the Department of Research and Innovation and the Department of Pathology, Nuvance Health for their help in patient recruitment.

## Supporting Information

**S1 Table. Serology results of Lyme disease patients by the year**.

**S2 Table. List of pathogenic bacteria used for specificity testing of TaqMan assays**.

**S3 Table. Cycle threshold (Ct) values from Real-time PCR analysis of 2017 patient samples**.

